# IntelliFold-2: Surpassing AlphaFold 3 via Architectural Refinement and Structural Consistency

**DOI:** 10.64898/2026.02.09.704787

**Authors:** Lifeng Qiao, He Yan, Gary Liu, Gaoxing Guo, Siqi Sun

## Abstract

IntelliFold-2 is an open-source biomolecular structure prediction model that improves accuracy and robustness through architectural refinement and multiscale structural consistency. We introduce latent space scaling in Pairformer blocks, a principled atom-attention formulation with stochastic atomization, policy-guided optimization for diffusion sampling and difficulty-aware loss reweighting. On Foldbench, IntelliFold-2 improves performance in therapeutically relevant settings, with particularly strong gains for antibody-antigen interactions and protein-ligand co-folding relative to AlphaFold 3. We release three variants (Flash, v2, and Pro) to cover efficient fine-tuning through high-precision server-side inference.

## Introduction

Accurate prediction of biomolecular structure and interactions is central to mechanistic biology and therapeutic discovery, with increasing emphasis on challenging settings such as antibody-antigen interactions and protein-ligand co-folding. Recent advances, most notably AlphaFold 3 [1], represent a landmark step for the field, substantially expanding the scope and reliability of unified biomolecular structure and interaction modelling.

Following our initial release [2], we present **IntelliFold-2**, a major update that delivers substantially improved performance. Concretely, IntelliFold-2 (i) scales Pairformer latent capacity to improve expressivity and hardware efficiency, (ii) enforces more principled multiscale structural representations through revised atom attention and stochastic atomization, (iii) improves diffusion-time reliability via policy-guided (PPO) optimization, and (iv) rebalances supervision with difficulty-aware loss reweighting. These changes lead to strong empirical gains on Foldbench [3], with particularly improved accuracy in antibody-antigen docking and protein-ligand co-folding relative to AlphaFold 3 [1].

Three model variants are provided:

- **IntelliFold-2-Flash:** A fast, efficient model intended primarily for academic use and easy fine-tuning, featuring our updated data curation and multiscale structural representations with 12 standard Pairformer blocks.
- **IntelliFold-2:** Our most accurate open-source model, featuring 48 widened Pairformer blocks with latent space scaling.
- **IntelliFold-2-Pro:** Our server-side flagship model. In addition to all architectural improvements, it incorporates exclusive PPO-enhanced sampling and Difficulty-Aware Loss optimization for maximum precision.

### Benchmark Performance

This release emphasizes performance improvements in structurally challenging yet therapeutically relevant categories, with a focus on **antibody-antigen interactions** and **protein-ligand co-folding**.

Building on the capabilities of the initial release, IntelliFold-2 now establishes a new state-of-the-art by outperforming AlphaFold 3 in these critical categories.

**Figure 1.**
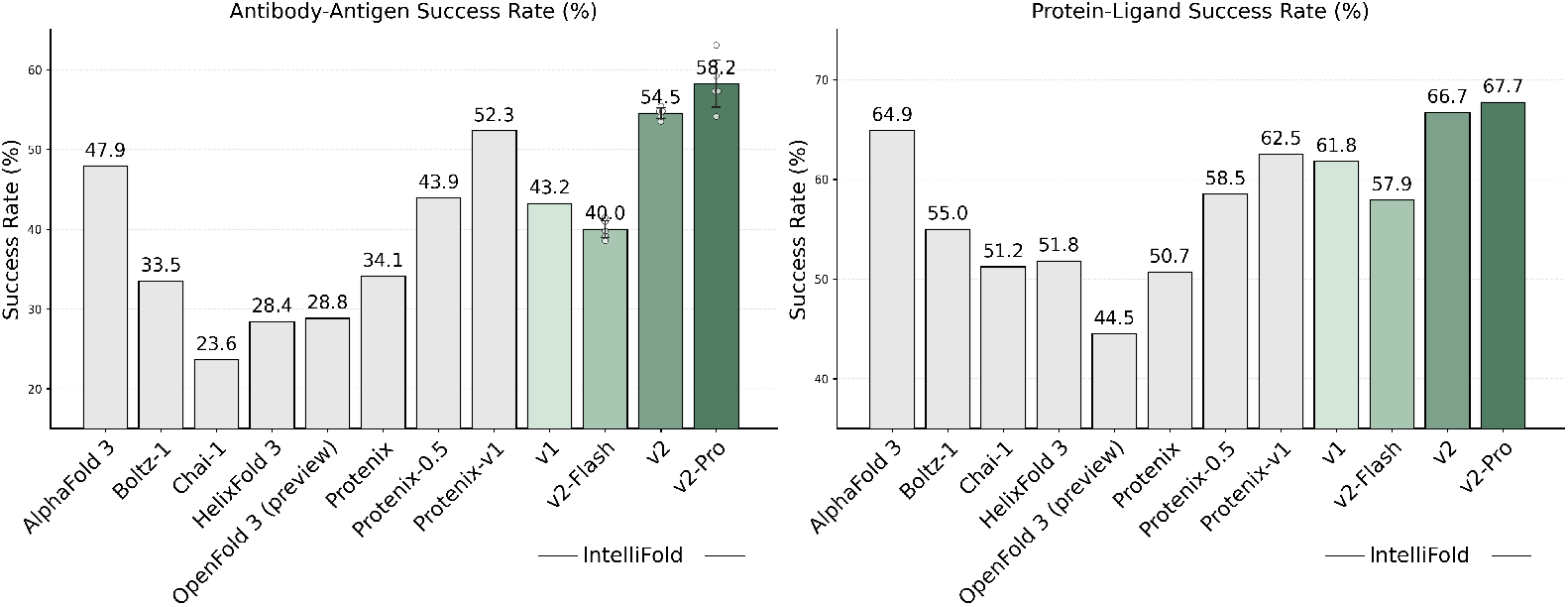
Performance on Foldbench [1, 2, 4–9]. IntelliFold-2 (v2 and v2-Pro) demonstrates a significant lead over AlphaFold 3 in Antibody-Antigen interactions and Protein-Ligand co-folding. Success is defined as DockQ > 0.23 for Antibody-Antigen and lRMSD < 2Å with LDDT-PLI > 0.8 for Protein-Ligand interactions. For ABAG, the reported v2-model results are aggregated over 5 runs, with raw datapoints provided.

### Key Technical Changes

#### Scaling Model Capacity and Computational Efficiency

Empirical analysis revealed that the previous hidden dimension bottlenecked both representational capacity and computational efficiency.

- **Latent Space Scaling:** We increase the dimensionality of latent representations within the Pairformer blocks, significantly enhancing the model’s capacity to capture complex biological interactions. Similar strategies were also explored at [10].
- **Improved Hardware Utilization:** Larger hidden dimensions increase arithmetic intensity, leading to substantially higher Model FLOPs Utilisation (MFU) and more effective use of modern GPU architectures.

#### Principled Multiscale Structural Representations

We refine the training procedure to better align atomic-level accuracy with global structural consistency.

- **Consistent Representations Across Scales:** We revise the atom attention mechanism to a more principled formulation, ensuring robust and self-consistent model behaviour during training and inference.
- **Stochastic Atomization:** Atom-level tokenization is applied stochastically to improve robustness and strengthen the model’s ability to capture fine-grained atomic interactions.

#### Policy-Guided Optimization for Diffusion Sampling

We apply reinforcement learning, instantiated via Proximal Policy Optimization (PPO), to fine-tune the diffusion module and improve sampling reliability at inference time.

- **Policy-Guided Sampling:** We frame the diffusion sampler as a stochastic policy and apply PPO updates to encourage trajectories that yield structurally coherent and physically plausible conformations.
- **Reduction of Random Sampling Failures:** By constraining policy updates through the clipped PPO objective, the fine-tuning process suppresses unstable or low-quality trajectories, leading to more robust sampling and fewer random failures during inference.

**Figure 2.**
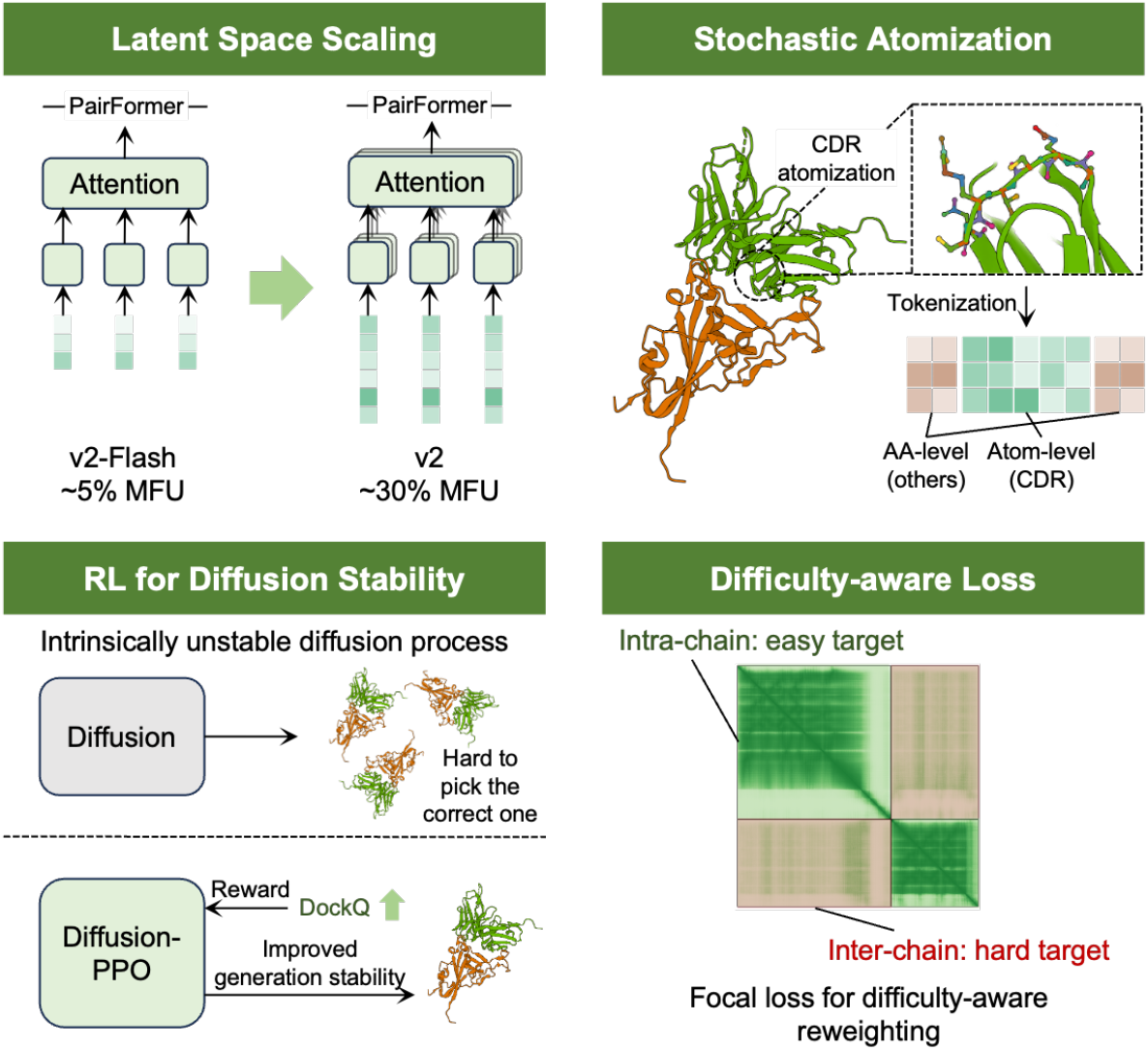
Key architectural and algorithmic innovations in IntelliFold-2

#### Difficulty-Aware Loss Reweighting

To address imbalance in training signals across structural difficulty levels, we adopt a focal-loss–style reweighting formulation.

- **Emphasis on Hard Examples:** Focal loss down-weights well-predicted samples and allocates more gradient signal to challenging regions of the structural space, such as flexible loops and ambiguous side-chain configurations.
- **Improved Optimization Dynamics:** This re-weighting leads to more stable convergence and improves accuracy on structurally complex targets without disproportionately affecting easy cases.

#### High-Fidelity and Scalable Data Curation

We re-processed the Protein Data Bank (PDB) dataset using an updated pipeline to ensure high-fidelity supervisory signals. Furthermore, we scaled up the self-distillation dataset and optimised its distribution. This expansion enriches the diversity of training examples and enhances the model’s generalisation capabilities across complex targets.

## Usage and Availability

IntelliFold-2-Flash and IntelliFold-2 are available at https://github.com/IntelliGen-AI/IntelliFold.

IntelliFold-2-Pro is available to Pro users via our online server at https://server.intfold.com.

## Conclusion

IntelliFold-2 advances open-source biomolecular structure prediction by combining scalable model capacity with self-consistent multiscale representations, more reliable diffusion sampling and difficulty-aware loss reweighting. These changes yield strong empirical gains on Foldbench, especially for antibody-antigen and protein-ligand tasks, while providing model variants that balance efficiency and accuracy.

## Author Contributions

**Lifeng Qiao** developed the majority of the technical changes. **He Yan** refined the atom local attention and validated the impact of self-distillation. **Gary Liu** and **Gaoxing Guo** contributed to building the distillation datasets. **Siqi Sun** led the project.

**Table 1:** Comparative performance metrics across diverse biomolecular systems. We report LDDT for monomers and % DockQ > 0.23 for interaction systems.

| Model | Protein Monomer | RNA Monomer | Protein-Protein | Protein-RNA |
| --- | --- | --- | --- | --- |
| AlphaFold 3 | 0.88 | 0.61 | 72.9 | 62.3 |
| IntelliFold-1 | 0.88 | 0.63 | 72.9 | 58.9 |
| IntelliFold-2-Flash | 0.88 | 0.55 | 73.6 | 56.5 |
| IntelliFold-2 | 0.89 | 0.58 | 71.9 | 68.3 |

## References

[1] Josh Abramson, Jonas Adler, Jack Dunger, Richard Evans, Tim Green, Alexander Pritzel, Olaf Ronneberger, Lindsay Willmore, Andrew J Ballard, Joshua Bambrick, et al. Accurate structure prediction of biomolecular interactions with alphafold 3. Nature, 630(8016):493–500, 2024.

[2] The IntFold Team, Leon Qiao, Wayne Bai, He Yan, Gary Liu, Nova Xi, Xiang Zhang, and Siqi Sun. Intfold: A controllable foundation model for general and specialized biomolecular structure prediction. arXiv preprint arXiv:2507.02025, 2025.

[3] Sheng Xu, Qiantai Feng, Lifeng Qiao, Hao Wu, Tao Shen, Yu Cheng, Shuangjia Zheng, and Siqi Sun. Benchmarking all-atom biomolecular structure prediction with foldbench. Nature Communications, 2025.

[4] Jeremy Wohlwend, Gabriele Corso, Saro Passaro, Noah Getz, Mateo Reveiz, Ken Leidal, Wojtek Swiderski, Liam Atkinson, Tally Portnoi, Itamar Chinn, et al. Boltz-1 democratizing biomolecular interaction modeling. BioRxiv, pages 2024–11, 2025.

[5] Chai Discovery team, Jacques Boitreaud, Jack Dent, Matthew McPartlon, Joshua Meier, Vinicius Reis, Alex Rogozhonikov, and Kevin Wu. Chai-1: Decoding the molecular interactions of life. BioRxiv, pages 2024–10, 2024.

[6] Lihang Liu, Shanzhuo Zhang, Yang Xue, Xianbin Ye, Kunrui Zhu, Yuxin Li, Yang Liu, Jie Gao, Wenlai Zhao, Hongkun Yu, et al. Technical report of helixfold3 for biomolecular structure prediction. arXiv preprint arXiv:2408.16975, 2024.

[7] The OpenFold3 Team. Openfold3-preview, 2025. URL https://github.com/aqlaboratory/openfold-3.

[8] ByteDance AML AI4Science Team, Xinshi Chen, Yuxuan Zhang, Chan Lu, Wenzhi Ma, Jiaqi Guan, Chengyue Gong, Jincai Yang, Hanyu Zhang, Ke Zhang, et al. Protenix-advancing structure prediction through a comprehensive alphafold3 reproduction. BioRxiv, pages 2025–01, 2025.

[9] Protenix Team, Yuxuan Zhang, Chengyue Gong, Hanyu Zhang, Wenzhi Ma, Zhenyu Liu, Xinshi Chen, Jiaqi Guan, Lan Wang, and Wenzhi Xiao. Protenix-v1: Toward high-accuracy open-source biomolecular structure prediction. bioRxiv, 2026. doi: 10.64898/2026.02.05.703733. URL https://www.biorxiv.org/content/10.64898/2026.02.05.703733v1.

[10] Yi Zhou, Chan Lu, Yiming Ma, Wei Qu, Fei Ye, Kexin Zhang, Lan Wang, Minrui Gui, and Quanquan Gu. Seedfold: Scaling biomolecular structure prediction, 2025. URL https://arxiv.org/abs/2512.24354.

